# Is there magnocellular facilitation of early neural processes underlying visual word recognition? Evidence from masked repetition priming with ERPs

**DOI:** 10.1101/2021.10.10.463440

**Authors:** Xin Huang, Wai Leung Wong, Chun-Yu Tse, Werner Sommer, Olaf Dimigen, Urs Maurer

**Affiliations:** Department of Psychology, The Chinese University of Hong Kong, Hong Kong, China; Department of Social and Behavioural Sciences, City University of Hong Kong, Hong Kong, China; Institut für Psychologie, Humboldt-Universität zu Berlin, Germany, Department of Psychology, Zhejiang Normal University, Jin Hua, China; Brain and Mind Institute, The Chinese University of Hong Kong, Hong Kong, China

**Keywords:** magnocellular, parvocellular, spatial frequency, luminance contrast, visual word, reading, EEG, ERP

## Abstract

An influential theory in the field of visual object recognition proposes that fast magnocellular (M) information facilitates neural processing of spatially more fine-grained but slower parvocellular (P) information. While written words can be considered as a special type of visual objects, it is unknown whether magnocellular facilitation also plays a role in reading. We used a masked priming paradigm that has been shown to result in neural facilitation in visual word processing and tested whether these facilitating effects are mediated by the magnocellular system. In two experiments, we manipulated the influence of magnocellular and parvocellular systems on visual processing of a contextually predictable target character by contrasting high versus low spatial frequency and luminance versus color contrast, respectively. In addition, unchanged (normal) primes were included in both experiments as a manipulation check. As expected, unchanged primes elicited typical repetition effects in the N1, N250 and P3 components of the ERP in both experiments. In the experiment manipulating spatial contrast, we obtained repetition effects only for the N1 component for both M- and P-biased primes. In the luminance versus color contrast experiment, repetition effects were found in N1 and N250 for both M- and P- biased primes. Furthermore, no interactions were found between M- vs. P-biased prime types and repetition. Together these results indicate that M- and P- information contributes jointly to early neural processes underlying visual word recognition.

## 1. Introduction

Reading has become an essential skill for human communication in everyday life. Compared with speaking and listening, reading can occur at faster rates with around 250 words per minute on average (see meta-analysis by Brysbaert, 2019). Such rapid processing of visual words is possible, as the visual system may speed up processing by making use of information that arrives through different visual pathways at different speeds, as has been suggested for object recognition (Bar, 2004). The human visual system is composed of three anatomically distinct parallel pathways (magnocellular M, parvocellular P, and koniocellular K) projecting from the magno-, parvo- and koniocellular ganglion cells to the lateral geniculate nucleus, and from there to the primary visual cortex V1. These cells have distinct structural and spatiotemporal characteristics. M cells are large cells with more myelinated axons which are sensitive to achromatic low spatial frequencies, have high contrast sensitivity and temporal resolution, and high conduction velocity (Kaplan, 2004). In contrast, P cells are small with less myelinated axons, are color-opponent (so-called red/green), and have low contrast sensitivity, high spatial and low temporal resolution, and a low conduction velocity (Yoonessi and Yoonessi, 2011). The M projections comprise most of the dorsal, or “where”, visual system, whereas the P projections form much of the ventral, or “what”, visual stream (Goodale and Milner, 1992). Some studies have found that the contrast sensitivity for color is shifted to lower spatial frequencies relative to the function for luminance (e.g., De Valois and De Valois, 1988). Besides, some psychophysical studies in humans indicate that the parvocellular system can mediate spatial contrast detection over a large range of spatial and temporal frequencies (Legge, 1978). Therefore, the magnocellular system may be more selectively activated by luminance changes than spatial frequency manipulations (Skottun, 2015).

Based on these physiological properties of the visual system, a model was proposed by Bar (2004) that can explain fast top-down facilitation in visual object recognition as part of a predictive coding framework. According to this model, the magnocellular system rapidly projects to orbitofrontal cortex, where “gist” information about the stimulus is extracted, which then facilitates subsequent occipito-temporal (OT) processing through top-down predictions. The proposed model is based on the findings that stimuli designed to bias processing toward M pathways differentially activated the OT compared with P-biased stimuli, and led to faster recognition and more accurate responses (Kveraga et al., 2007).

OT has been found to be associated with not only visual object recognition but also visual word recognition (Dehaene and Cohen, 2011; Price and Devlin, 2011). During visual word recognition, the ventral occipito-temporal (vOT) region is consistently engaged, and damage to this area will cause reading deficits (e.g., alexia; Starrfelt et al., 2009). Besides, vOT also acts as an interface associating bottom-up visual form information critical for orthographic processing with top-down higher order linguistic properties of the stimuli in some neural models (Kherif et al., 2011; Nakamura et al., 2005). The importance of the vOT region for both object and visual word processing raises the question of whether similar mechanisms that have been proposed for object recognition may also facilitate visual word recognition.

In fact, written words may be considered as a kind of visual object (Schlaggar and McCandliss, 2007), and they also contain spatial frequency and luminance contrast information. Normally, low spatial frequencies (LSF) provide word-shape information, such as length, general shape and location (Boden and Giaschi, 2009), whereas high spatial frequencies (HSF) provide fine-grained features like oriented lines and edges, which are crucial for letter identification (Majaj et al., 2002). There is no doubt that word recognition in reading relies on HSF input (Beckmann et al., 1991; Majaj et al., 2002). However, an important role of LSF information in visual word recognition cannot be ignored. Some studies have found that word shapes, and perhaps by extension LSF information, can help word identification. For example, when isolated (Beech and Mayall, 2005), the outer word fragments of a word can provide more relevant information for lexical access than the inner ones. Besides, words presented with the features of the global word form available could facilitate lexical access (Underwood and Bargh, 1982). Some studies also found that when letters were low-pass filtered by (e.g., at 1.1 cycles per letter, Kwon and Legge, 2012), letter recognition accuracy remained close to 100%. All the evidence above indicates that readers can extract global, coarse information to facilitate the recognition of letters or words.

Similar to spatial frequency, color and luminance contrast can also influence reading. Legge et al. (1990) found that luminance and color contrast influenced reading independently as no additive interaction of luminance and color was found, suggesting that the neural signals used for letter recognition are carried by both pathways for color and luminance. Besides, there is consistent evidence that the M pathway is involved in processing text. For example, with the allocation of attention, magnocellular input could help flanked-letter identification (Omtzigt and Hendriks, 2004).

In addition, some studies on dyslexia suggest that the magnocellular system is involved in visual word recognition. Several studies found that dyslexic readers have lower contrast sensitivity than controls (Demb et al., 1998; Felmingham and Jakobson, 1995; Lovegrove et al., 1980; but see Brannan and Williams, 1988; refer to Skottun, 2000 for a recent review), or have deficits in the magnocellular system or dorsal-stream visual pathway (Livingstone et al., 1991). And the proposal that dyslexia is the result of a magnocellular deficit in the visual system has gained some support (Boets et al., 2011; Talcott et al., 2000). Taken together, several lines of evidence suggest that magnocellular information is utilized in visual word recognition.

One experimental paradigm that has provided facilitation effects in visual word recognition is the masked priming paradigm. In this paradigm, a prime is presented for a very short duration followed (and often preceded) by a mask (backward masking), which makes the prime consciously imperceptible. However, even with a brief (and unconscious) exposure, the prime can produce robust effects on subsequent processing, as indicated by shorter reaction times and lower error rates for targets after valid primes compared to targets after invalid primes. One of the classical findings is a repetition effect, in which, when the prime and target are the same word, reaction times to the target words are shorter compared to the baseline condition when the prime and target are different, unrelated words (see, Forster, 1998, for a review). More recently, several studies used event-related potentials (ERPs) in the EEG to investigate neural mechanisms involved in visual word processing (Eddy et al., 2006; Grainger et al., 2012; Holcomb et al., 2005; Holcomb and Grainger, 2006; Kiefer and Brendel, 2006). In alphabetic languages, masked repetition of visual words led to a reduction in the N250 and N400 components of the ERP at central electrodes (with mastoid reference) compared to unrelated words (Chauncey et al., 2008; Holcomb and Grainger, 2006). The N250 repetition effects seem to be sensitive to visual-orthographic properties of the words, as it can be modulated when primes and targets show orthographic overlap. The reduction of the N250 and N400 components may be considered as a correlate of facilitation of orthographic and semantic processes (Grainger et al., 2012; Holcomb and Grainger, 2006). In addition to the N250 and N400, the P3 was also found to be sensitive to masked priming manipulation, with a topographic maximum over posterior regions (Holcomb and Grainger, 2006). The P3 was more positive in the repeated condition compared to unrelated primes, presumably reflecting the priming the whole-word representation, as it was modulated by a complete overlap between prime and target words.

In addition to the N250, N400, and P3 effects, some studies also found repetition effects in the N1 (N/P 150) component with increased negativity for same versus different word presentations at occipital electrodes (Chauncey et al., 2008). This N1 effect seems to be less consistent and less robust compared to the later effects in these studies, possibly due to the use of a mastoid reference that is close to the electrodes where the N1 is maximal. The N1 component has been shown to be sensitive to print, as it was larger for letter strings in familiar writing systems compared to visual control stimuli or word forms from unfamiliar writing systems (Maurer et al., 2005, 2008). Recent studies with more closely matched control stimuli suggest that this print tuning effect mainly manifests in the N1 onset, possibly reflecting an earlier onset of print-specialized processing (Eberhard-Moscicka et al., 2016; Wang and Maurer, 2017, 2020). In a model of the time-course of neural correlates of visual word recognition that was based on EEG masked priming studies, the N1 component was suggested to reflect the mapping of visual features to location-specific letter positions in alphabetic languages (Grainger and Holcomb, 2009).

Masked repetition priming studies in Chinese showed a similar result pattern as those in alphabetic languages. Parallel to studies in alphabetic languages, repeated targets elicited a repetition priming effect with reduced amplitudes of N250 (Wong et al., 2014) and N400 (Du et al., 2014) compared to different word primes. No masked priming effects were reported in these studies for the N1 component. These studies show that masked priming effects in early components are commonly obtained in visual word recognition irrespective of the properties of the writing system, and that the N250 repetition effect might be a good candidate to test facilitation at the level of early visual orthographic processing across different writing systems.

Some studies have already investigated the role of LSF information in masked repetition priming. A behavioral study conducted by Boden and Giaschi (2009) used a lower band of LSF (2 cycles per degree, cpd) and a higher band of HSF (8 cpd) stimuli and found that neither HSF nor LSF information was sufficient to prime lexical decisions. When they changed the bandwidth of LSF to higher cut-off frequencies of 3.5 cpd and lower HSF of 4.6 cpd, they found that both HSF and LSF information could produce equivalent priming effects, indicating that LSF was unlikely to have a unique role in word recognition. However, it is noteworthy that like many other masked priming studies, Boden and Giaschi (2009) presented isolated prime-target pairs, meaning that words were not expected or predictable from the context. In contrast, natural reading situations typically afford at least some predictions about the upcoming word and it is conceivable that top-down processing mediated via the magnocellular pathway may be more relevant under these conditions.

A recent study by Winsler et al. (2017) first used ERPs to investigate whether both HSF and LSF information contributes to visual word recognition. Using a masked priming paradigm, the authors presented HSF (above 15.2 cpd), LSF (below 3.7 cpd) and FSF (full spatial frequency) primes and measured the N1, N250 and N400 components. For the HSF and FSF conditions, repetition effects were significant for both the N250 and N400 components, but not for the N1 component. Besides, the LSF primes elicited a distinct early repetition effect around 200 ms (N250) that was opposite in direction to typical repetition effects, indicating that coarse information played a role in visual word recognition. However, the inversed polarity of the effect might better correspond to a N1 effect (Chauncey et al., 2008) although the timing is late. Given that the FSF condition also showed a similar but nonsignificant pattern, and that the reference electrode used was not very sensitive to the occipital N1 effect, further investigation is warranted. Furthermore, as in the Boden and Giaschi (2009) study, the lack of expectational inducement, as it is typical for natural reading situations, may have obscured any magnocellular effects on N250 facilitation in this study. In addition, as mentioned before, it is possible that luminance and color contrast are more effective manipulations of magnocellular processing as compared to spatial frequency. Given these limitations, it seems justified to further examine whether magnocellular information influences N250 facilitation in a masked priming experiment that allows for predictions about the upcoming word and also uses a luminance manipulation.

Furthermore, we conducted our study in Chinese, which potentially is more sensitive to visual and M-biased manipulations. On average, Chinese characters have more strokes than letters, which may put more demands on visual processing (McBride-Chang et al., 2011; Zhao et al., 2014). Also, complex characters usually require higher contrast thresholds (Yu and Cao, 1992) and higher spatial frequencies (Wang and Legge, 2018) for identification compared with simple characters. Besides, Chinese characters are densely packed, hence, they may contain more configural visual information (which defines spatial relations between components or radicals, e.g., up-down, right-left) than linear alphabetic letters. There is consensus that the stroke sequences contain HSF information, whereas the configural relations among components depend on LSF information (Perfetti et al., 2013). Therefore, LSF information may be more useful for Chinese than alphabetic reading.

To induce expectations, we used Chinese two-character words as materials and presented them sequentially character-by-character, which should help the reader to form an expectation for the upcoming characters. Chinese words consist of one or more characters. Compound words, especially two-character words, account for a dominant proportion of words in Chinese. In two-character words, both characters can be morphemes. Thus, when the first character is presented, it can help activate morpheme family words (Wang and Liu, 2019). To further induce expectations, we selected only two words of the morpheme family by matching some variables (see Methods below) to control the conditional probability among a word pair. Therefore, when the first character was presented, the readers’ expectations on the second character would be similar. In traditional masked priming studies, researchers usually used word pairs as prime and target (e.g., chair–table in English, 国王–皇帝 [king–empire] in Chinese). In this case, words are processed as a holistic unit, with little predictability of the following word based on the first word. In the current study, we presented Chinese two-character words in a character-by-character way, and the prime was either identical or unrelated to the second character of the words, hence readers can form expectations on the upcoming characters.

To investigate the relative contributions of the magnocellular and parvocellular pathways in visual word recognition, we manipulated the content of primes through manipulating the spatial frequency (Experiment 1) and luminance contrast (Experiment 2). As in Winsler et al. (2017), three manipulations on spatial frequency were used in Experiment 1, they were HSF, LSF, and unchanged (FSF). The logic of using the manipulations was that FSF primes should replicate the typical masked repetition effects, HSF primes should trigger parvocellular processing, and LSF primes should tap into magnocellular processing. We followed the same manipulations as in Boden and Giaschi’s study (2009). To create HSF primes, we high-pass filtered at 8 cpd like some previous studies (Carretié et al., 2020; Vaegan and Hollows, 2006). For LSF primes, we note that Legge (1978) suggested 1.5 cpd as the cross-over point. However, since we used a masked priming paradigm, which may diminish the impact of LSF information, a low pass filter below 2 cpd was used in the current study, which is also in line with some previous studies (Felmingham and Jakobson, 1995; Lovegrove et al., 1980).

Experiment 2 manipulated color and luminance contrast in order to extend the findings from Experiment 1. The design was similar to Experiment 1, except that we manipulated the content of primes by using luminance contrast and color in unchanged, isoluminant, and heteroluminant conditions. Given the characteristics of the magnocellular and parvocellular systems, the masked priming effects should be stronger under the manipulation of luminance contrast.

By combining ERPs with masked priming repetition effects, we hypothesized that, if there was an interaction between repetition and spatial frequencies/luminance contrast, it would indicate that different spatial frequencies/luminance contrasts may play different roles during visual word recognition. The unchanged conditions in the two experiments served as a manipulation check: Here, we expected an attenuation of the N250 and N400 components and an increased N1 and P3 for targets preceded by repeated primes as compared to unrelated primes. For M- and P-biased manipulations, we expected N250 repetition priming effects for M-biased primes (LSF/heteroluminant) rather than for P-biased primes (HSF/isoluminant), therefore showing a magnocellular facilitation in early neural processes of visual word recognition.

## 2. Methods

### 2.1. Participants

Twenty-four native Mandarin Chinese participants (17 females; mean age = 21.06 years old, range = 21–29) took part in the two experiments during the same session. All participants reported to have normal vision or corrected-to-normal visual acuity and normal color vision, reported having neither dyslexia nor ADHD and were right-handed as determined by the Chinese Handedness Questionnaire (Li, 1983). Written informed consent was obtained before the experiment. All participants were reimbursed 50 Hong Kong dollars (about 7 USD) per hour. The study was approved by the Joint Chinese University of Hong Kong-New Territories East Cluster Clinical Research Ethics Committee.

### 2.2. Materials

Forty pairs of two-character words were selected from the SUBTLEX-CH database of Chinese word and character frequencies (Cai and Brysbaert, 2010). Among these word pairs, the first character was the same in both words (for example, **搏**斗 – **搏**击). Pairs were made so that the second character of the other word of the pair could serve as the unrelated prime in the unrelated priming condition for the other word. All words were of medium or high frequency in the language. As shown in Table 1, the two words of each pair fairly closely matched according to word frequency, word contextual diversity^1^ (CD, Adelman et al., 2006), character frequency, character contextual diversity, number of strokes, logographeme number of the second character. There was no shared radical in the second character with the first character among a word pair. All 80 two-character words (40 pairs) were presented together with identical and unrelated primes in three priming conditions (M-biased, P-biased, and unchanged primes) resulting in 480 critical trials. An additional 72 target words were randomly selected from the 480 critical trials and added to the list as probes, for which participants needed to name the word (proportion of probe trials: 13%). This resulted in a total number of 552 trials in each of the two experiments.

**Table 1.** Mean differences of the characteristics between the words in the pair (standard deviation in parentheses).

| Characteristics | Stroke number | Number of Logographs | Character frequency | Character CD | Word frequency | Word CD |
| --- | --- | --- | --- | --- | --- | --- |
| Mean difference | 0.48 (1.41) | 0.13 (1.41) | 0.01 (1.47) | 0.01 (1.17) | 0.002 (0.05) | 0.01 (0.18) |

Note. The first four characteristics are extracted from the second character of a word, and the last two characteristics are extracted from the whole word. Frequency and CD values are log-transformed.

The first character and target stimuli were presented in black within a small square (3 cm × 3 cm) with a white background (except the M- and P-biased manipulation in Experiment 2, see below) presented in Songti font. Primes were instead presented in a Kaiti font to reduce visual overlap between prime and target stimuli. Stimuli were viewed from a distance of about 70 cm so that primes and targets subtend 2.68° of horizontal and vertical visual angle. In Experiment 1, the M- and P-biased primes were spatially filtered (using function *fft* in Matlab) thereby leaving only high (> 8 cpd; P-biased) or only low spatial frequencies (< 2 cpd; M-biased). In Experiment 2, achromatic, heteroluminant stimuli were used as M-biased primes; P-biased primes were chromatic with the character and its background being isoluminant (red characters shown on a green background with the same luminance, see Fig. 1B). Heteroluminant stimuli had a mean Michelson contrast^2^ of 20% (Green et al., 2009). To make sure the luminance of the colors of the background and characters in the P-biased conditions were the same, we measured the luminance values with a Minolta LS-150 photometer. In addition, an unchanged condition was included in both experiments to replicate typical masked priming effects.

**Fig. 1.**
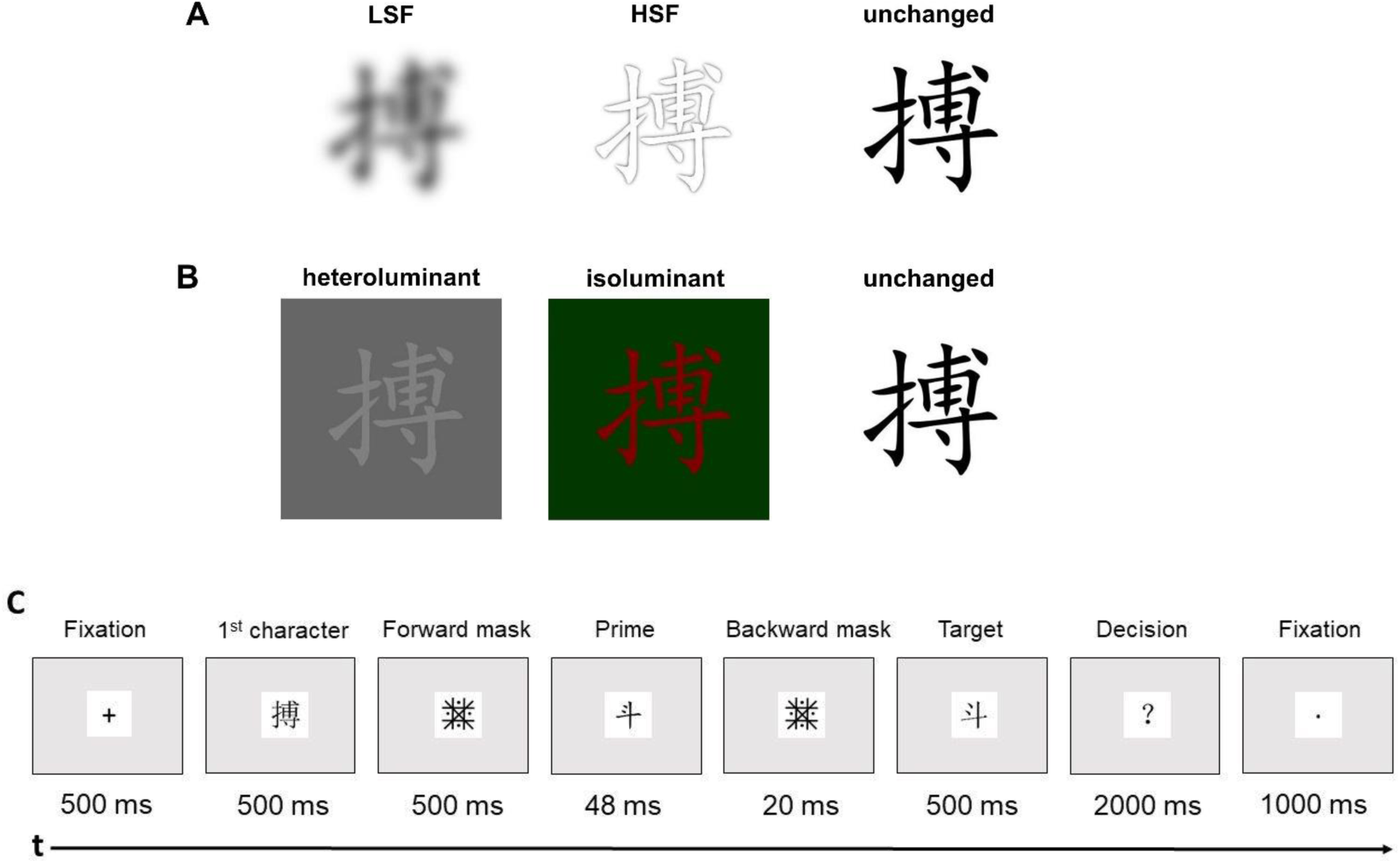
Examples of primes in Experiment 1 **(A)** and Experiment 2 (**B**) with a schematic illustration of a typical trial (**C)**. Non-probe trials were shown without the 2000 ms question mark slide.

### 2.3. Procedure

Participants were seated in a comfortable chair, viewing a computer monitor with a refresh rate set at 144 Hz in a sound-attenuated darkened room. The testing session began with a short practice block, followed by one of the six counterbalanced experimental blocks. Participants performed a naming task in which they were instructed to verbally repeat the character whenever the target character was followed by a question mark (probe). Each session comprised six blocks, with two blocks each of unchanged, P-biased, and M-biased conditions. Participants could rest during breaks between blocks. The six experimental blocks in each experiment were counterbalanced by using a Latin square design such that every target would occur in both the repeated and unrelated conditions and in each spatial frequency/luminance contrast condition. The order of items within each block was pseudorandomized. The order of the two experiments was also counterbalanced.

As shown in Fig. 1C, each trial began with a fixation cross in the middle of the screen; after 500 ms, the first character was presented. A forward mask, consisting of intersecting vertical, horizontal, and diagonal lines (Perfetti and Zhang, 1991) that occupied the same space as the characters was presented for 500 ms. The forward mask was then replaced at the same location on the screen by a prime in Kaiti font for 48 ms, consisting of the second character of the other word (unrelated) or the same word (repeated) in the pair. The prime was then immediately replaced by a backward mask. The backward mask remained on screen for 20 ms and was then immediately replaced by the target character in Songti font for a duration of 500 ms. In the trials that included probes, the target was followed by a question mark for 2 s, indicating that participants needed to name the second character of a word. Each trial ended with a 1-s fixation point. Participants were asked to refrain from blinking and moving their eyes except when the fixation point was on the screen.

After the two EEG experiments, we did a debriefing about how the experiment was composed to participants, and they were asked whether they noticed the primes during the experiments. Afterward, participants completed a writing task, in which they needed to write down the primes. The procedure of the writing task was similar to the formal experiment, except that only unchanged primes were presented to the participants, and participants had no time limit to write down the prime characters they saw. The accuracy of the writing task was calculated for each participant.

### 2.4. EEG recordings

The EEG was recorded from 62 Ag/AgCl scalp electrodes mounted in a textile cap at standard 10–10 positions and referenced against CPz. Two electro-oculogram (EOG) electrodes were placed on the outer canthus of each eye and one EOG was placed on the infraorbital ridge of the left eye. Signals were amplified with an ANT amplifier system (Advanced Neuro Technology, Enschede, Netherlands) at a bandpass of 0.01–70 Hz and sampled at 1000 Hz. Impedances were kept below 20 kΩ.

### 2.5. EEG pre-processing

EEG data were analyzed using Brain Vision Analyzer (Brain Products GmbH, Gilching, Munich). The continuous EEG was re-referenced to average reference (Lehmann and Skrandies, 1980)) and digitally filtered using IIR band pass Butterworth filter with a 0.3–30 Hz (24 dB/octave) bandpass. EEG signals were decomposed into maximally-independent processes via extended-infomax Independent Component Analysis (ICA, Jung et al., 2000), and ocular-related components were identified through visual inspection and then removed. Before averaging, the trials were segmented from 150 ms prior to 850 ms after the stimulus onset. Trials with amplitudes exceeding ±80 µV in any channel were automatically rejected from further analyses. All participants had similar numbers of trials for all conditions in the two experiments (see Table 2 for the trial number of each condition in each experiment). After baseline correction (–150–0 ms), grand means were computed for all conditions.

**Table 2.**
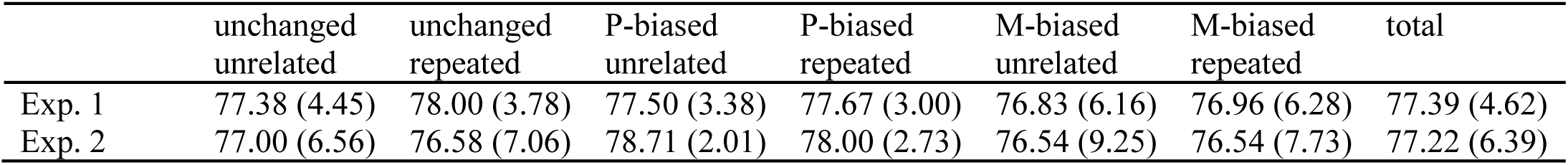
Mean trial numbers and standard deviations (in parentheses).

### 2.6. Data analysis

Separate ERPs were calculated for the six types of target conditions in each experiment. The analysis strategy was to first compare targets in the repeated and unrelated prime conditions with unchanged stimuli to identify time windows with repetition effects, and then use those time windows to test whether repetition effects would differ between M- and P-biased stimuli. To identify the time windows of repetition effects in unchanged stimuli, we ran point-to-point Topographic Analyses of Variance (TANOVA; Koenig et al., 2011) on non-normalized (raw) scalp maps comparing target ERPs following valid primes to those following invalid primes for unchanged conditions, separately for the two experiments. The TANOVA was corrected for multiple comparisons through Global Duration Statistics (Koenig et al., 2011). Based on the TANOVA results, we selected the time windows in which repetition effects were significant (*p* < .05). As we were mainly interested in repetition effects corresponding to the N1 and N250 effects, we focused on effects within the first 300 ms after stimulus presentation. Then we applied the time windows to M- and P-biased conditions to average amplitudes of regions of interest (ROI). The selection of ROI was based on the t-maps in which electrodes showed the maximal activation in the unchanged contrast. The t-maps were computed by subtracting the repeated condition from the unrelated condition in Brain Vision Analyzer based on the time windows identified by TANOVA (see Fig. 4).

Repeated measures analyses of variance (ANOVAs) were performed on these measurements with the factors Repetition (Repeated vs. Unrelated), Laterality (Left vs. Right), and Type (LSF vs. HSF in Experiment 1, heteroluminant vs. isoluminant in Experiment 2). The Greenhouse-Geisser correction was applied to all effects with more than one degree of freedom in the numerator (Greenhouse and Geisser, 1959).

## 3. Results

Visual inspection of the target ERP waveforms at OT electrodes showed an early positive peak around 40 ms after the onset of the target stimulus and an early negative peak around 90 ms followed by a broad subsequent negativity with additional negative peaks at around 140 and 230 ms (see Fig. 2). Given their early latencies, the first positive and negative peaks presumably reflect a P1-N1 complex elicited by the preceding prime (presented at –68 ms), while the later negative peaks presumably reflect the N1 and N250 components in response to the targets. As on average the prime and target stimuli were the same in the repeated and unrelated conditions, contrasting the two conditions will result in differences that are elicited by the relationship between the prime and target only (see, e.g., Grainger and Holcomb, 2009).

**Fig. 2.**
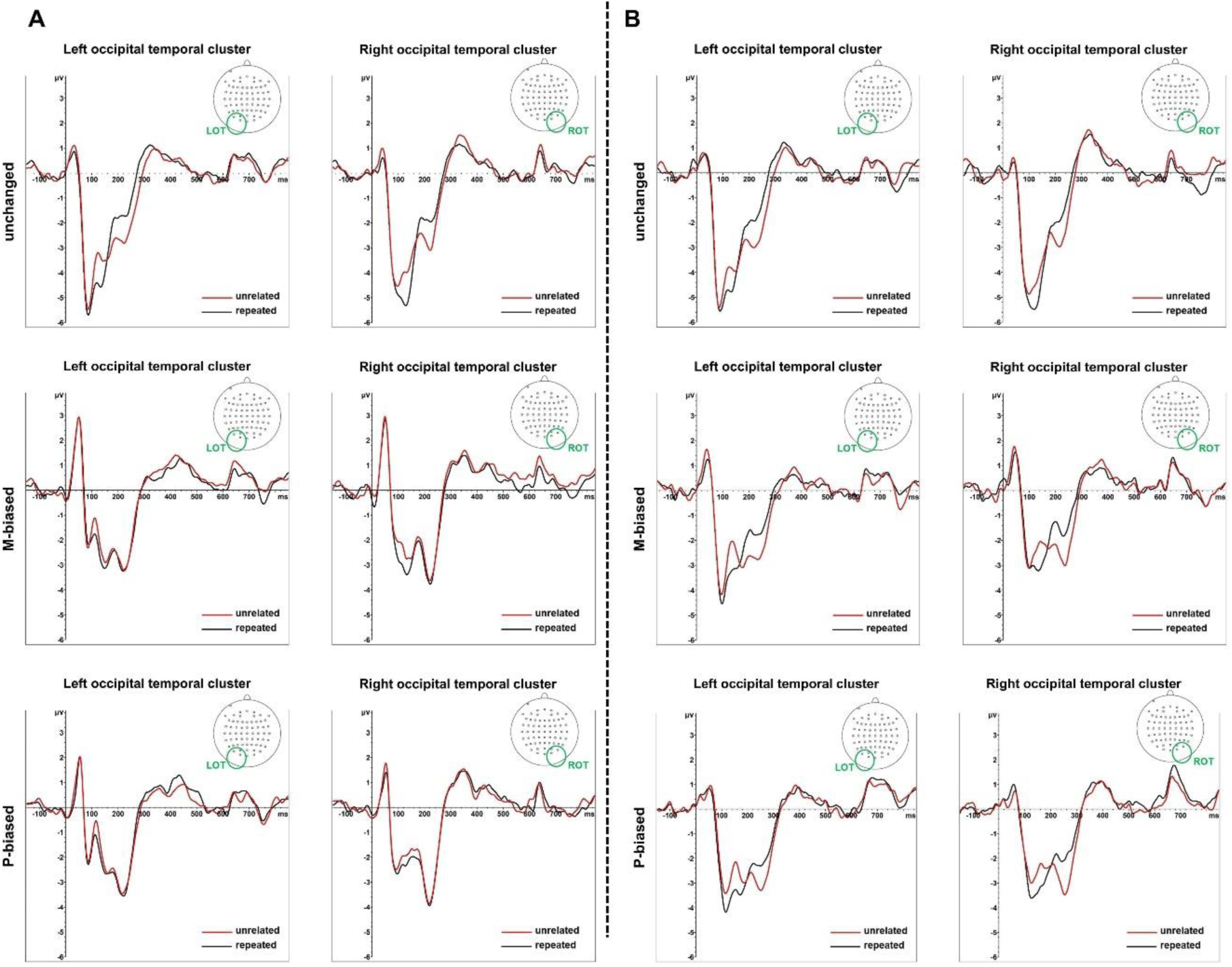
Comparison of target-ERP waveforms as a function of prime type (unchanged, M-biased, P-biased). ***A,*** the ERP waveforms at left occipito-temporal (LOT) and right occipito-temporal (ROT) electrode clusters for repeated and unrelated primes in unchanged (top), HSF (middle), and LSF primes (bottom) of Experiment 1. ***B,*** the ERP waveforms at left occipito-temporal (LOT) and right occipito-temporal (ROT) electrode clusters for repeated and unrelated primes in unchanged (top), isoluminant (middle), and heteroluminant primes (bottom) of Experiment 2.

### 3.1. EEG results

#### 3.1.1 Experiment 1 – spatial frequency manipulation

TANOVA comparing repeated and unrelated targets within the unchanged prime condition revealed two time windows. The first window lasted from 103–163 ms, consistent with a repetition priming effect in the N1 component in the waveforms or a P1-N1 transition based on the timing in visual ERPs. The t-map indicated that occipito-temporal electrodes (PO7/PO8, P7/P8, O1/O2) showed a maximal difference (see Fig. 4A). A significant N250 repetition effect was found in two time windows that were separated by merely two sampling points (see Fig. 3A). As the *p*-value of these two time points was below 0.1, we combined them for further analysis, resulting in a larger time window from 195–296 ms. Similar to the N1, OT electrodes (PO7/ PO8, P7/P8, O1/O2) showed the maximal difference for this N250 effect. For the P3 component, the time window was measured in a 405–453 ms segment, and the central regions (CP1/CP2, CP3/CP4, CP5/CP6, P1/P2, P3/P4) showed maximal activation according to t-maps. In addition, a 5^th^ time window was identified between 596–653 ms ^3^ (see Fig. 3A for TANOVA results).

**Fig. 3.**
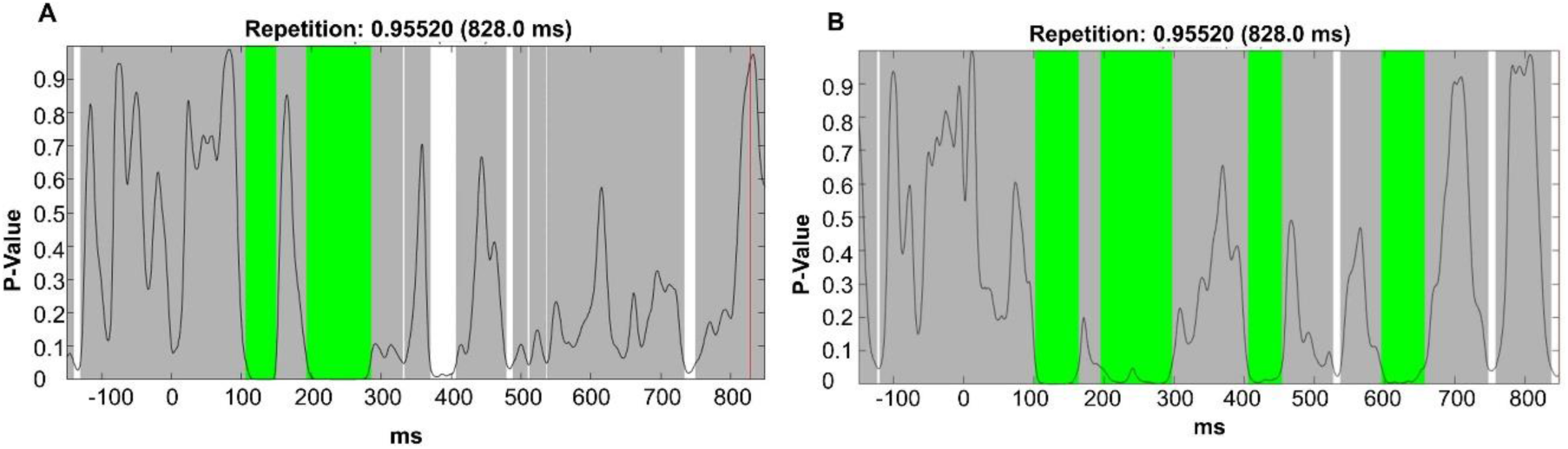
Results of sample-by-sample TANOVAs with global duration statistics (marked in green) in Experiment 1(**A**) and Experiment 2 (**B**). For Experiment 1, the duration threshold was identified as 43 ms, the threshold was then applied to the TANOVA plot, where periods longer than the estimated duration threshold are marked in green. For Experiment 2, the duration threshold was identified as 41 ms.

**Fig. 4.**
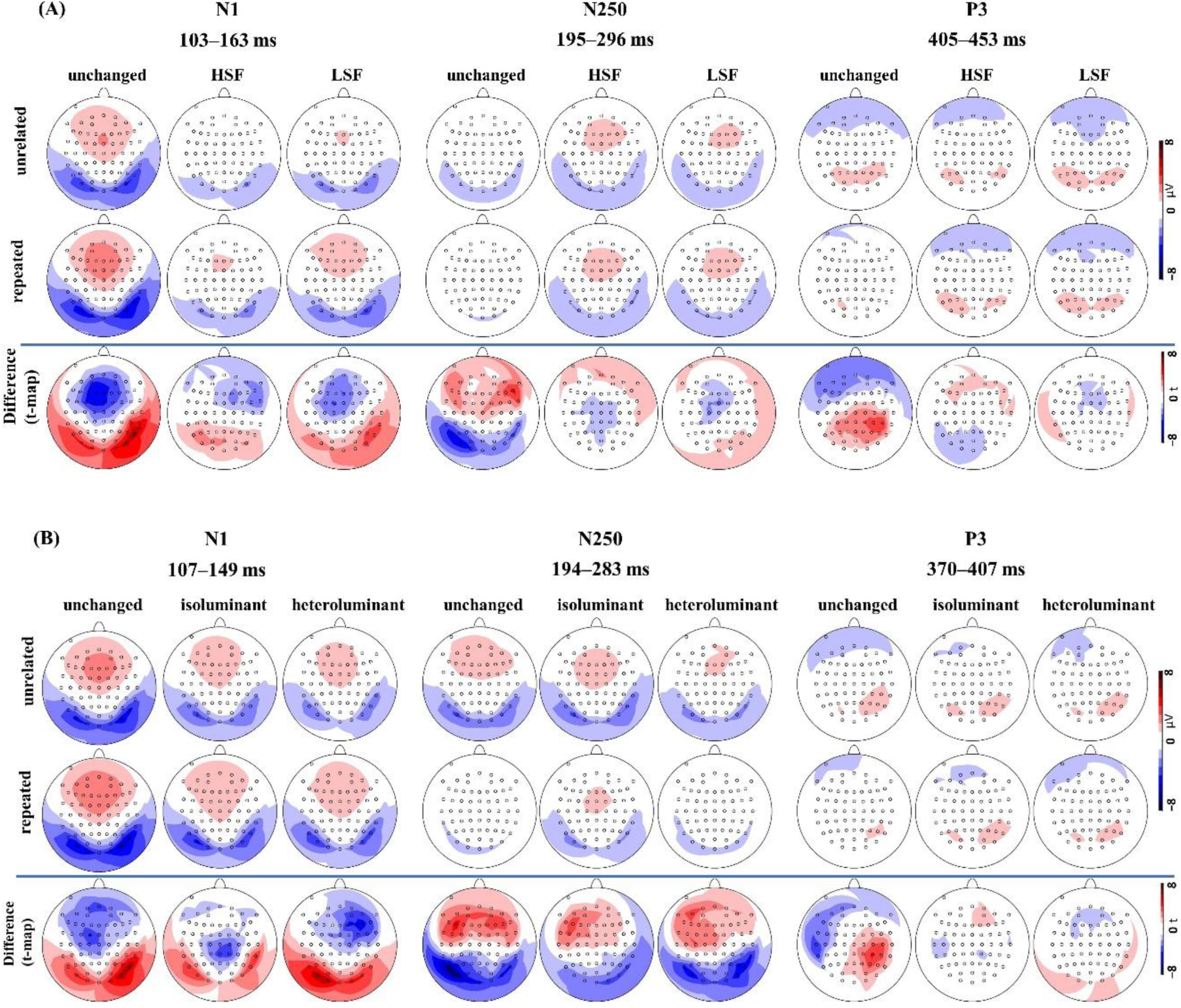
***A,*** ERP effect topographies and t-maps for the repetition priming effect according to prime type. Data is shown at the N1 (103–163 ms), N250 (195–296 ms), and P3 (405–453 ms) in Experiment 1. ***B,*** the ERP and t-maps at the N1 (107–149 ms), N250 (194–283 ms), and P3 (370–407 ms) for each type of prime in Experiment 2.

##### 3.1.1.1 N1 (103–163 ms)

To examine whether LSF had a greater repetition effect than HSF, we ran a three-way ANOVA with factors Type, Repetition, and Laterality. The results revealed that the N1 was increased for repeated compared to unrelated targets (*Repetition*, *F* _(1, 23)_ =12.15, *p* = 0.002, 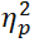 = 0.35, see Fig. 3A). However, the repetition effect was not different between the LSF and HSF primes (*Repetition* × *Type*, *F* _(1,23)_ = 0.50, *p* = 0.49, 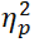 = 0.02). In addition, regardless of prime relation, N1 amplitude to the target was larger following LSF than HSF primes (*Type*, *F* _(1, 23)_ = 9.00, *p* = 0.006, 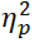 = 0.28). Neither the main effect of lateralization nor any other interactions were significant (*F*s < 0.80, *p*s > 0.38). Follow-up *t*-tests on the average of left and right OT regions showed that repetition effects in HSF (*t* _(23)_ = 2.22, *p* = 0.036, *d* = 0.45) and LSF (*t* _(23)_ = 3.06, *p* = 0.006, *d* = 0.63) were both significant, with repeated characters showing larger activation than unrelated characters.

To test whether the results at ROI reflected the effects across entire ERP maps, TANOVA (not normalized) on average amplitudes in the N1 time window was computed with factor type (HSF vs. LSF) and repetition (repeated vs. unrelated). The results showed a significant repetition effect (*Repetition*, *p* = 0.02) that did not differ between HSF and LSF conditions (*Repetition × Type*, *p* = 0.70). In addition, the ERP maps differed between HSF and LSF conditions irrespective of prime relation (*Type*, *p* = 0.001). The distribution of the repetition effects is illustrated in t-maps (see Fig.4A).

##### 3.1.1.2. N250 (195–296 ms)

Similar to the analysis of the early N1, we ran three-way ANOVA for the LSF and HSF conditions. Neither the main effect of type, repetition and laterality nor the interactions were significant (*F*s < 1.75, *p*s > 0.20). Follow-up *t*-tests on the average of left and right OT showed that repetition effects in HSF (*t*_(23)_ = 0.17, *p* = 0.87) and LSF (*t*_(23)_ = 0.83, *p* = 0.42) were not significant.

TANOVA results showed no significant repetition effects across two types of primes (*Type*, *p* = 0.301; *Repetition*, *p* = 0.443; *Repetition × Type*, *p* = 0.846).

##### 3.1.1.3. P3 (405–453 ms)

We first ran three-way ANOVAs with factor Type, Repetition and Laterality. Results showed that neither the main effects nor the interactions were significant (*F*s < 2.21, *p*s > 0.15). Follow-up *t*-test on the average of central regions showed that repetition effects in HSF (*t* _(23)_ = 0.81, *p* = 0.43) and LSF (*t* _(23)_ = 0.64, *p* = 0.53) were not significant.

Supplementary TANOVA results showed no significant repetition effects across two types of primes (*Repetition*, *p* = 0.825; *Repetition × Type*, *p* = 0.614), although ERP maps differed between HSF and LSF conditions irrespective of prime relation (*Type*, *p* = 0.038).

#### 3.1.2. Experiment 2 – luminance manipulation

Based on the principles outlined in the methods section, we first report the results of TANOVA and which segments and electrodes were used for the subsequent analyses. For the early N1 component, the time window was identified as 107–149 ms. The 194–283 ms time window was identified for the N250 component. For the P3 component, the time window was measured in a 370–407 ms epoch. The selection of ROI in N1 and N250 was identical to Experiment 1, whereas the t-maps showed the central area (P5/P6, CP1/CP2, CP3/CP4, CP5/CP6, FC3/FC4, FC5/FC6) to have maximal activation in P3, which was slightly different from Experiment 1.

##### 3.1.2.1. N1 (107–149 ms)

To examine whether the heteroluminant condition had a greater repetition effect than isoluminant conditions, we ran a three-way ANOVA with factors Type, Repetition, and Laterality. The results revealed that the N1 was increased for repeated compared to unrelated targets (*Repetition*, *F* _(1,23)_ = 40.26, *p* < 0.001, 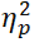 = 0.64). However, neither the interaction (*Type× Repetition*, *F* _(1,23)_ = 0.24, *p* = 0.63, 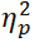 = 0.01) nor the main effect of type *(F* _(1,23)_ = 1.66, *p* = 0.21, 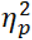 = 0.07) was significant, indicating that heteroluminant and isoluminant conditions did not produce different patterns in N1. Follow-up *t*-test on the average of left and right OT regions showed that repetition effects in isoluminant (*t* _(23)_ = 3.96, *p* = 0.001, *d* = 0.81) and heteroluminant conditions (*t* _(23)_ = 3.73, *p* = 0.001, *d* = 1.15) were both significant, with repeated characters showing larger N1 amplitudes than unrelated characters (separate comparisons in each condition are shown in Fig. 3B).

In the N1, TANOVA showed a significant repetition effect (*Repetition*, *p* = 0.004) that did not differ between heteroluminant and isoluminant conditions (*Repetition × Type*, *p* = 0.352). ERP maps tended to differ between heteroluminant and isoluminant conditions irrespective of prime relations (*Type*, *p* = 0.057). the distribution of the repetition effects is illustrated with t-maps (see Fig. 4B).

##### 3.1.2.2. N250 (194 –283 ms)

We first ran the three-way ANOVA for isoluminant and heteroluminant conditions. The results revealed that the N250 amplitude was reduced for repeated compared to unrelated targets (*Repetition*, *F* _(1,23)_ = 20.84, *p* < 0.001, 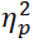 = 0.48). And isoluminant primes led to larger N250 negativity in the subsequent target than heteroluminant primes (*Type*, *F* _(1, 23)_ = 9.42, *p* = .005, 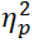 = 0.29). However, the repetition effects did not differ between heteroluminant and isoluminant conditions (*Type× Repetition, F* _(1, 23)_ = 0.91, *p* = 0.35, 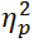 = 0.04). Follow-up *t*-test on the average of OT regions showed that repetition effects in isoluminant (*t* _(23)_ = –3.25, *p* = 0.004, *d* = 0.66) and heteroluminant (*t* _(23)_ = –4.71, *p* < 0.001, *d* = 0.96) conditions were both significant, with unrelated target characters showing larger N250 amplitudes than repeated characters.

TANOVA showed a significant repetition effect (*Repetition*, *p* = 0.028) that did not differ between heteroluminant and isoluminant conditions (*Repetition × Type*, *p* = 0.924). In addition, ERP maps differed between heteroluminant and isoluminant conditions irrespective of prime relations (*Type*, *p* = 0.003).

##### 3.1.2.3. P3 (370–407 ms)

The three-way ANOVAs including factor Type (isoluminant vs. heteroluminant) showed that neither the main effects nor the interactions were significant (*F*s < 2.64, *p*s > 0.12). Follow-up *t*-test on the average of central regions showed that the repetition effect in the isoluminant condition (*t* _(23)_ = –1.79, *p* = 0.087, *d* = 0.37) was marginally significant, and the repetition effects in heteroluminant condition (*t*_(23)_ = –0.02, *p* = 0.987, *d* = 0.003) was not significant.

In the P3, we did not find significant repetition effects or interactions in TANOVA (*Type*, *p* = 0.713; *Repetition*, *p* = 0.922; *Repetition × Type*, *p* = 0.488).

### 3.2. Behaviour results

Naming accuracy for the target in the occasional probe trials was high in both experiments, with 98.4% (*SE* = 0.02, range: 90.3–100%) for Experiment 1 and 98.7% (*SE* = 0.02; range: 88.8–100%) for Experiment 2.

The analysis of the writing task after the EEG experiments showed that participants, on average, had an accuracy of 50.7% (*SE* = 0.285) in reporting the prime character; the accuracy for 18 of the participants was below 50%. To further test whether the repetition effects were influenced by the visibility of the primes, we correlated the accuracy in the writing task with the size of the repetition effects on the three ERP components of the two experiments (see Supplementary Table 1). We found that the accuracy in the writing task correlated with the size of the repetition effect in P3 of Experiment 2 (*r* = 0.409), but not with that in the other components.

## 4. General Discussion

The present study aimed to investigate whether magnocellular information would facilitate the early neural processes underlying visual word recognition when the expectation about upcoming stimuli is induced. As we recorded ERPs in a masked repetition priming paradigm and with Chinese characters, an additional aim was to determine whether neural facilitation would already occur at the level of the N1 component to Chinese characters, particularly when using an average reference. In addition to normal, unchanged primes, we presented P-biased (parvocellular) and M-biased (magnocellular) primes by separately manipulating spatial frequency (Experiment 1) and luminance versus color contrast (Experiment 2). To induce expectations, we used Chinese two-character words and presented them character-by-character, hence, participants could form an expectation about the full two-character word on the basis of the first character (prime).

For unchanged (normal) primes, we obtained repetition priming effects in both experiments that corresponded to the N1 (or P1-N1 transition), N250, and P3 components that were then used to analyze priming effects of M- and P-biased primes. Similarly, both P- and M-biased primes induced repetition priming effects with an increased N1 negativity (or faster P1-N1 transition). For the N250, P- and M-biased repetition effects were significant for the luminance manipulation but not for the spatial frequency manipulation. No significant repetition effects for M- or P-biased primes were found in the P3, neither for luminance contrast nor for the spatial frequency manipulation. Importantly, no interaction was found between repetition and the M-/P-based primes in either of the two experiments and in any of the ERP components analyzed. In other words, whether the prime was biased towards processing in the parvo- vs. magnocellular pathway did not affect the size of the repetition effect.

In both experiments, we obtained an early N1 repetition effect, with a scalp distribution that was more negative for repeated words at occipital sites and more positive at frontal sites. While the timing of this effect coincides with a broad negativity in the ERP with several small peaks, the early latency of this effect (of around 100 to 150 ms) may actually correspond to the P1-N1 transition in regular ERPs, where the P1 typically occurs at around 100 ms, and the N1 at around 150 ms. The repetition priming effect in this time window, therefore, would be in agreement with a forward-shift of the N1 component, although this latency shift may only be visible as an amplitude effect in masked priming experiments due to the overlapping ERP components elicited by the preceding mask and prime.

The direction of the repetition effect and the scalp distribution of the N1 effect was similar to that found in previous studies with repeated words, in which the size of the N/P150 was enhanced (Chauncey et al., 2008). While N1 repetition effects have not been consistently reported in previous masked repetition priming studies, the robust effects in the current study may be due to the use of an average reference that results in larger sensitivity for early visual components over occipital areas compared to the average mastoids reference (Lehmann and Skrandies, 1980). Another potential reason for the pronounced N1 repetition priming effect is that we used Chinese characters that are visually more complex than alphabetic letters, which puts higher demands on visual processing (Perfetti et al., 2013). Previously, it has been suggested that the N1 reflects rather basic processes in visual word recognition such as mapping visual features to letters (Holcomb and Grainger, 2007) or processing familiar features in letter strings (Maurer et al., 2005). A larger number of visual features may require more visual processing, which may render repetition effects in Chinese more robust than in alphabetic languages.

Like previous studies using alphabetic languages (Holcomb and Grainger, 2006; Petit et al., 2006), we obtained the typical repetition effect in the N250: targets that repeated across prime and target positions showed a reduced N250 negativity compared to those following unrelated primes. The direction of the current N250 repetition effect was the same as in previous studies (Holcomb and Grainger, 2007, 2006), although the N250 was only a negative-going component in the previous studies (Holcomb and Grainger, 2006, 2007). Moreover, the N250 effect in the previous studies was largest at parietal electrodes (e.g., Holcomb and Grainger, 2006, 2007), whereas it was largest at OT electrodes in the current study with a polarity reversal at frontal electrodes. The scalp distribution of the N250 effect in our study is similar to the distribution of N1/N250 components in previous studies on visual word processing that were localized to occipito-temporal regions (Brem et al., 2005; Maurer et al., 2005), consistent with fMRI findings showing that vOT is involved in visual word recognition (e.g., Kherif et al., 2011). The difference in the location of the effects can be explained by the use of an average reference in our study instead of a mastoid reference. Difference maps are always forced to zero at the location of the reference electrode; hence, a difference map with mastoid reference will always have very small amplitudes around the mastoid(s). In contrast, difference maps with average reference do not have a predetermined zero point. In addition, the average reference allows for a more straightforward interpretation suggesting that the N250 effect corresponds to reduced activation after repetition in agreement with a repetition suppression effect reported in fMRI studies (Henson et al., 2000).

Following the N250, we obtained P3 repetition effects in the unchanged conditions in both experiments with increased parietal positivity in response to targets following unrelated primes compared to targets following repetition primes. The P3 repetition effect in the current study resulted from increased parietal positivity of a positive component peaking at around 350–400 ms for unrelated targets compared to repeated targets (see Fig. 2 in the supplementary materials). The distribution of the P3 effect was similar to the distribution of the P3 and N400 effects in previous studies that were maximal at parietal electrodes, but the direction of the effect was the opposite (Holcomb and Grainger, 2006), which cannot be explained by the different reference. Instead, we think the presence of the P3 repetition effect in our study is a consequence of the particular paradigm used in our study in which a two-character Chinese word was presented sequentially in a character-by-character way. The P3 is thought to reflect the representations within the lexical level of processing (Grainger and Holcomb, 2009), and it might be sensitive to the outcome of the mismatch process described in the preceding context and its consequences for word-level processing. Due to the expectations induced in our experiment, a mismatch may be detected in the unrelated conditions that would engage additional processes to verify that lexical representations do indeed fit with the lower-level orthographic activation.

We observed N1 repetition effects not only for unchanged primes, but also for M- and P-biased primes in both experiments. While the study by Winsler et al. (2017) used masked repetition priming with a spatial filter manipulation, no N1 repetition priming effects were reported in that study presumably for the same reasons, as discussed above. The presence of the N1 repetition effects in our study across all different prime manipulations suggests that readers can benefit from partial visual information to process visual features.

A repetition priming effect in the N250 component, as it was found after unchanged primes, could only be found after primes with luminance or color contrast (in Experiment 2), but not after primes with spatial frequency contrast (in Experiment 1). This is different from the study by Winsler et al. (2017) who found a repetition priming effect for HSF stimuli, while an effect for LSF primes occurred earlier with a reversed direction. The different findings regarding N250 repetition effects with LSF primes may be explained by the writing system differences, as Chinese characters are visually more complex and may require more visual orthographic processing. Another potential reason for the differences between the two studies is that we induced the expectation on the upcoming characters so that readers could generate predictions about the visual form of the words (Gagl et al., 2020).

The presence or absence of the N250 repetition effects in the two types of magnocellular manipulation in our study may also be informative about the N250 repetition priming effect itself. As this effect only occurred with luminance and color contrast primes, but not with LSF and HSF primes, processes that occur during the N250 seem to benefit from the information that retains the original spatial frequency information of the stimulus, but may not benefit from partial spatial frequency information.

In the P3 component, only unchanged, but not M- and P-biased primes elicited repetition effects, indicating whole-word representation requires more than just LSF/HSF or luminance contrast information, fine-grained information is necessary for lexical access. We noted that Winsler et al. (2017) did not report any significant repetition effect in P3, in contrast, they observed a repetition effect for HSF with high-frequency words in N400. However, we did not obtain any repetition effect neither on N400 nor P3 for M-/P-biased primes^4^. Winsler et al. (2017) used a higher threshold of HSF than in our study (15.2 cpd vs. 8 cpd), thus more LSF information was retained in our study. And the different findings also suggest that lexical access depends on how much visual information is kept during visual word recognition, and the middle range of spatial frequency may not be beneficial for visual word recognition (i.e., between 8 to 15.2 cpd).

Taking the findings from the analyses of the repetition effects together, we found no evidence that the size of repetition effects differs for M- vs. P-biased primes in any of the components. This suggests that the repetition priming effects in our study were not mediated specifically by either the M system, as one might have expected based on models of object recognition (Bar, 2004). This is consistent with the findings of Winsler et al., (2017) with LSF stimuli. The current study corroborates and extends this null effect by showing that repetition priming effects do not differ between M-biased and P-biased primes when a different visual manipulation is used (luminance/color contrast) or when an inducement is put on the expectation of the upcoming stimulus. The results, therefore, do not support the view that magnocellular information from the same word is fed forward very fast and then provides top-down inducement for parvocellular information from the same word, as suggested for object recognition (Bar, 2004). In contrast, our results suggest that both P-biased and M-biased visual information is necessary to obtain the full range of facilitation effects, as seen with unchanged stimuli. However, our study does not rule out that M-biased information plays an important role in reading at all. The stimuli in our experiment were all presented in the center of the screen and were supposedly processed by the foveal region of the retina. It is possible that magnocellular information plays a role in reading, but mainly if it comes from the upcoming word that is processed in the parafoveal and peripheral regions of the visual field during natural reading. In fact, P-cells are well represented in and around the fovea, but M-cells are better represented in the retinal periphery (Kaplan, 2004), which may make the repetition effects more pronounced when words are presented in the parafovea. Therefore, further studies may address the issue of potential magnocellular facilitation in the parafoveal area, for example by combining EEG with eye-tracking.

We are aware of several potential limitations of the current study. Firsts of all, it is possible that the manipulation of the M system co-activates the other visual system, although the thresholds of our experiments were based on previous studies (Boden and Giaschi, 2009; Kveraga et al., 2007). For example, the contrast gain at a higher contrast (above 30%) of M cells is similar to that of the P cells, so M cells still respond to contrast modulation at high contrast. And P cells outnumber M cells and could compensate for their lower contrast gain by converging on cortical neurons (Kaplan, 2014). Therefore, it is possible that the activation of one visual system is accompanied by the other visual system. Even though we may have not exclusively involved one system over the other one, it is still likely that with our paradigm we achieved a relative dominance of processing within one system.

In addition, we noted that the M- and P- biased stimuli, which are not commonly encountered during natural reading, were less familiar to participants compared to the unchanged primes. The disruption of the processing stream could therefore also be a consequence of stimulus familiarity. To minimize this possibility, we used a masked priming paradigm such that the primes could be unconsciously perceived. And we also used different fonts for primes and targets to reduce the visual familiarity. To control the visibility of the primes, we used a pattern mask in our study, which is more effective than other previously used masks (such as the hash mask (#), Schoonbaert and Grainger, 2004)) as it is more visually complex. Furthermore, we correlated the repetition effects with the accuracy in the writing task (refer to the Table 1 in supplementary materials), and we did not find any significant correlations except a moderate positive correlation between P3 in Experiment 2 and the accuracy. The results indicated that early repetition effects were not influenced by the prime visibility. The moderate correlation for the P3 of Experiment 2 indicates that a larger repetition effect may be associated with higher prime visibility, therefore, the prime visibility may only influence the late phase of visual word recognition.

To conclude, our results suggest that neural facilitation of visual-orthographic processes in masked repetition priming of words not only occurs via a reduced N250 activation, but also via an increased or potentially accelerated N1 activation. Moreover, neural facilitation effects of visual word processing by masked primes seem to be neither mediated by M-biased information nor by P-biased information alone. While either type of information seems able to elicit facilitation of early visual processing, a combination of both types of information appears to be necessary to lead to facilitation at higher (presumably semantic) levels of cognitive processing.

## Acknowledgements

This work was supported by the General Research Fund of the Research Grants Council of Hong Kong (RGC-GRF 14616418).

## Supplementary materials

**Table 1.** Pearson correlation coefficients of the repetition effects in N1, N250 and P3 in the two experiments and the accuracy in the writing task

|  | N1 (Exp. 1) | N250 (Exp. 1) | P3 (Exp. 1) | N1 (Exp. 2) | N250 (Exp. 2) | P3 (Exp. 2) | ACC |
| --- | --- | --- | --- | --- | --- | --- | --- |
| N1 (Exp.1) | 1 | .697** | 0.215 | .945** | .677** | 0.178 | -0.069 |
| N250 (Exp.1) |  | 1 | .415* | .687** | .948** | .588** | 0.251 |
| P3 (Exp. 1) |  |  | 1 | 0.088 | 0.289 | .786** | 0.34 |
| N1 (Exp. 2) |  |  |  | 1 | .693** | 0.155 | -0.1 |
| N250 (Exp. 2) |  |  |  |  | 1 | .518** | 0.246 |
| P3 (Exp.2) |  |  |  |  |  | 1 | **.409\*** |
| ACC |  |  |  |  |  |  | 1 |

\*\* . Correlation is significant at the 0.01 level (2-tailed).
\* . Correlation is significant at the 0.05 level (2-tailed).
Note. ACC means the accuracy in the writing task.

**Supplementary Fig. 1.**
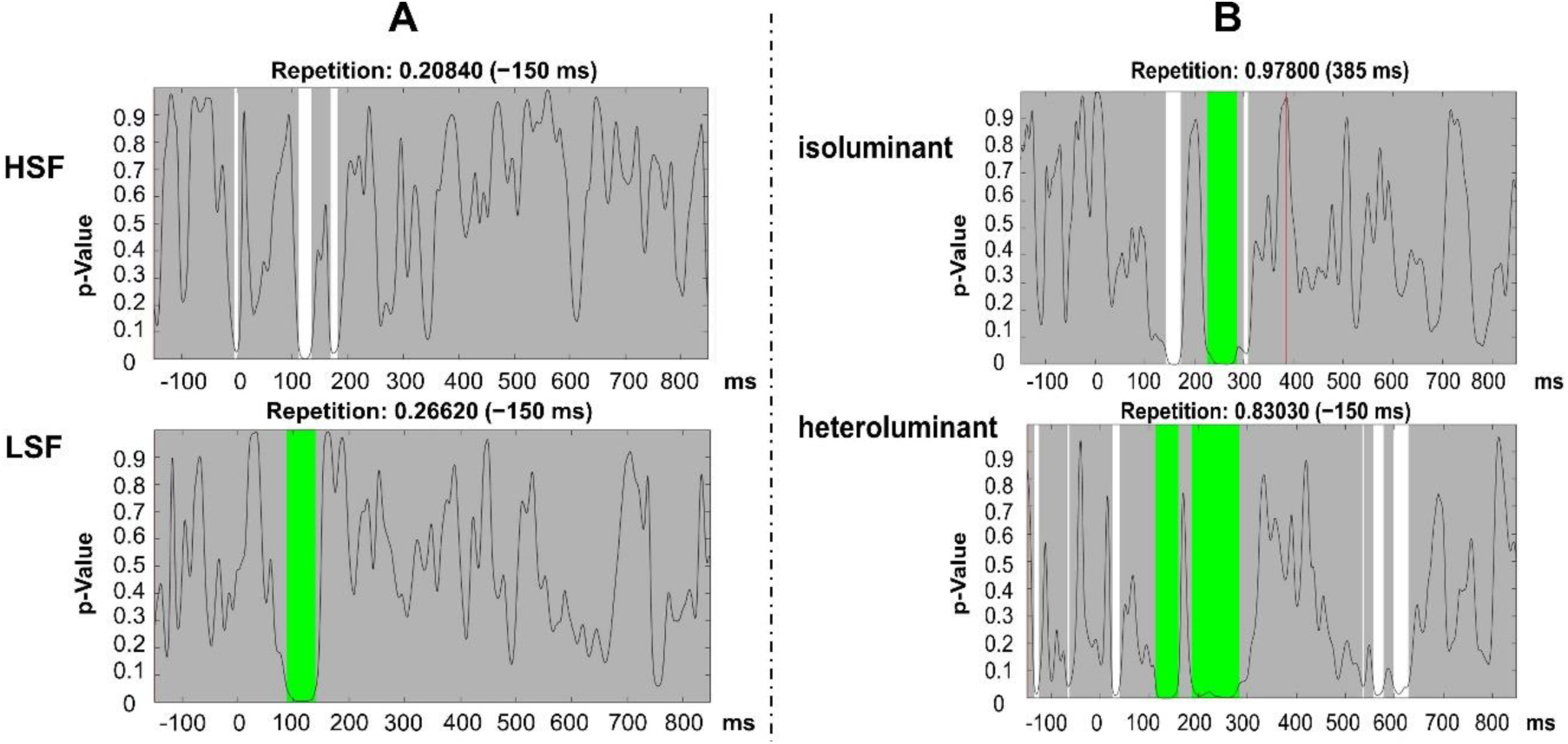
Point-to-point TANOVA comparing repeated and unrelated targets on the HSF and LSF conditions for Experiment 1 (**A**) and isolumiant and heterolumiant for Experiment 2 (**B**). The duration thresholds identified from global duration statistics for HSF, LSF, isoluminant and heteroluminant condition were 44 ms, 44 ms, 45 ms, 44 ms respectively. The duration thresholds were applied to TANOVA plots, where periods longer than the estimated duration threshold are marked in green.

**Supplementary Fig. 2.**
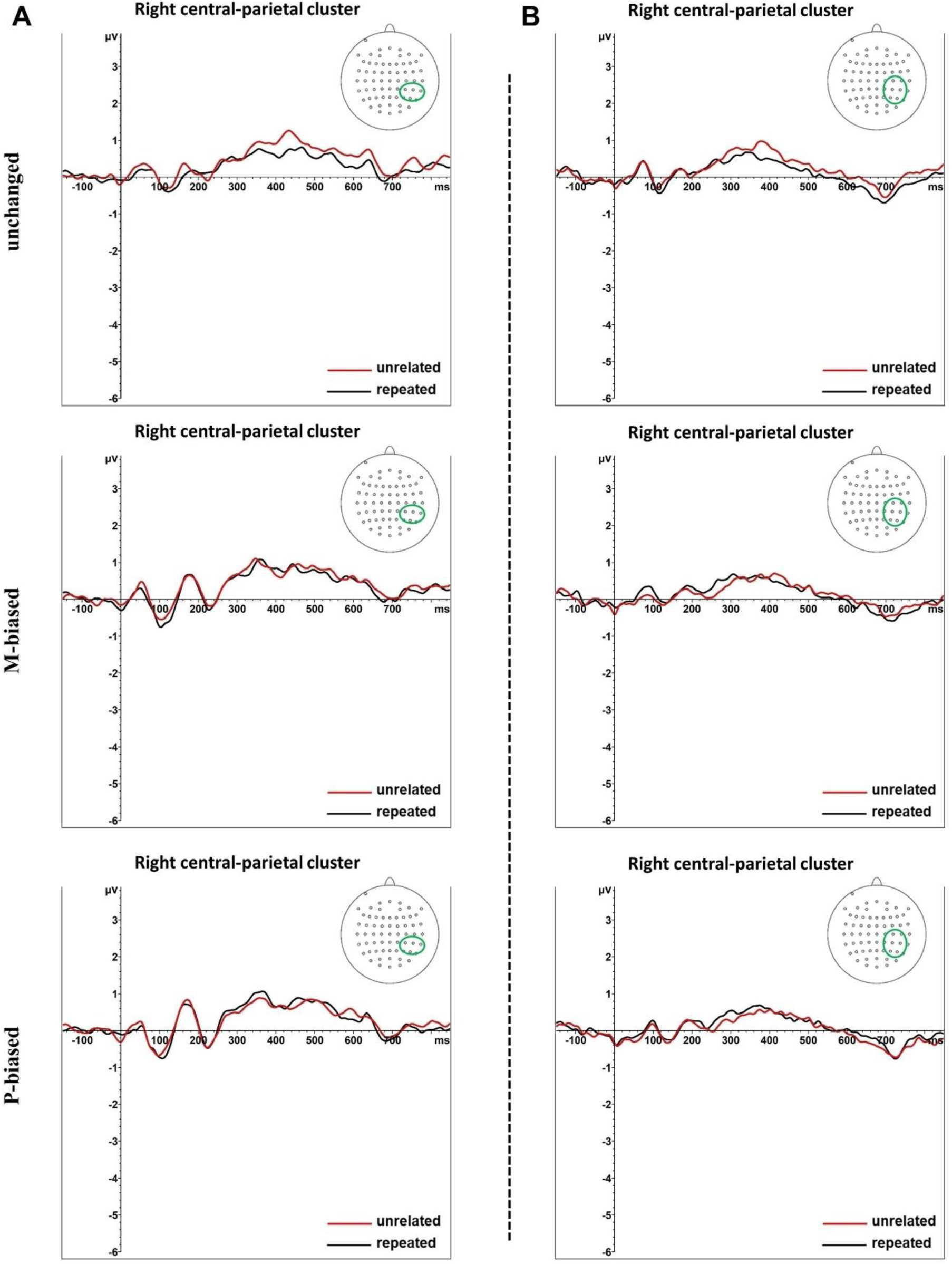
Separate ERP waveform comparisons for each prime type. The ERP waveforms at right central-parietal electrode clusters for repeated and unrelated primes in unchanged (top), M-biased (middle), and P-biased primes (bottom) of Experiment 1 (left panel) and Experiment 2 (right panel).

## Footnotes

1 Contextual diversity is defined as the proportion of texts in which a given word occurs.

2 Michelson contrast is defined as the absolute difference of fore- and background luminance values divided by their sum.

3 We noted that TANOVA identified a late time window in Experiment 1, which is not identified in Experiment 2. To investigate the different findings in the late phase, we ran a repeated ANOVA with a covariate of temporal order of experiment. The repetition effect was not significant (*F* _(1,22)_ = 0.034, *p* = 0.85).

4 We noted that the time windows of P350 identified by TANOVA were slightly different, and the time window of the P350 in Experiment 2 did not survive after correction, suggesting that the P350 effects are more variable in time. Therefore, to further test whether we could find any repetition effects in P350 in different time windows, we ran a point-to-point TANOVA comparing repeated and unrelated targets on the M- and P-biased primes for each experiment (see Supplementary Fig. 1). The TANOVA results showed that for M/P- biased primes, no other time windows after 300 ms of stimulus onset was identified except for heteroluminant condition, two late time windows which were separated by 19 ms were identified. By combining the two separate time windows, we obtained a time window of 557–628 ms. Together with the waveforms and the timing, it was unlikely the later effect reflecting P350.

